## Supplementary material for "Endothelin B receptor inhibition rescues aging-dependent neuronal regenerative decline": Supp Material and Figures

**S5:** KEGG pathway analysis for SGC is aged vs adult DRG

#### SUPPLEMENTARY TABLES

**Supplementary Table 1:** list of qPCR Primers

#### SUPPLEMENTARY MOVIES

**Movie S1:** Z-stack video of whole mount preparation of DRG from Lycopersicon Esculentum Lectin (LEL) injected Fabp7CreER::Ai14 mouse, which labels SGCs with tdTomato, immunostained for TUJ1 (neurons).

**Movie S2:** Z-stack video of DRG section immunostained for FABP7 and CX43.

### Figure 1- Figure Supplement 1

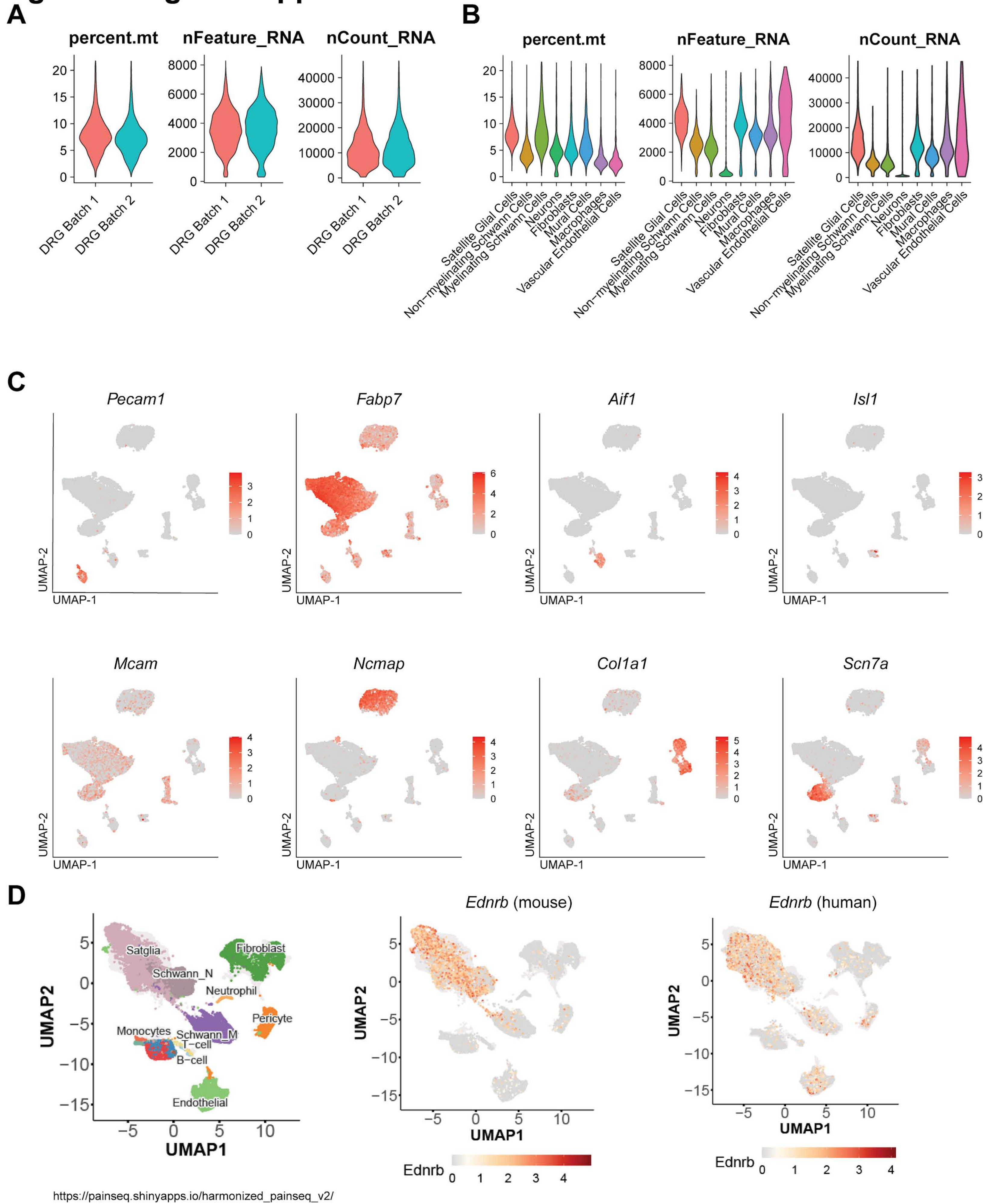

### Figure 3- Figure Supplement 1

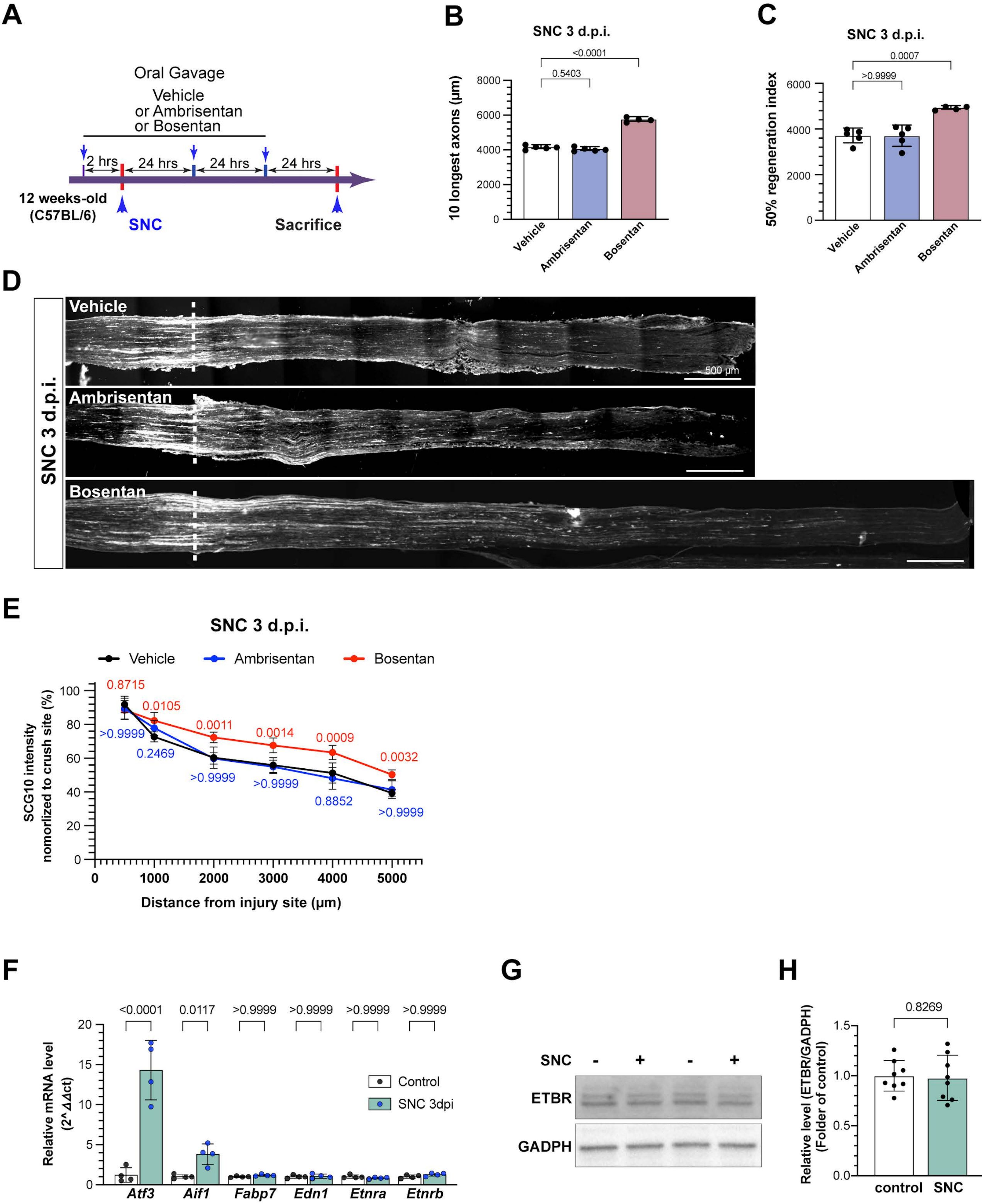

### Figure 3- Figure Supplement 2

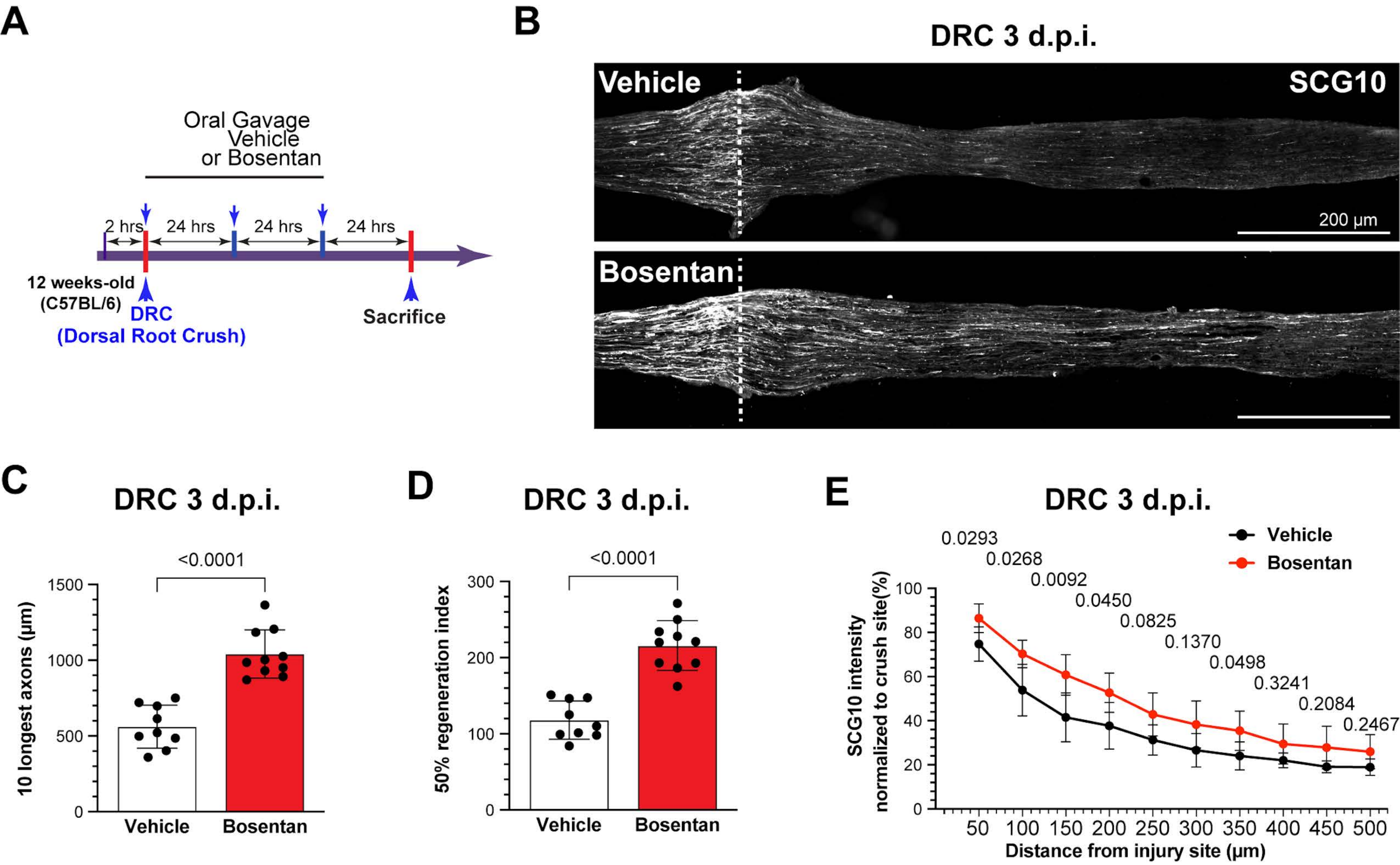

### Figure 4- Figure Supplement 1

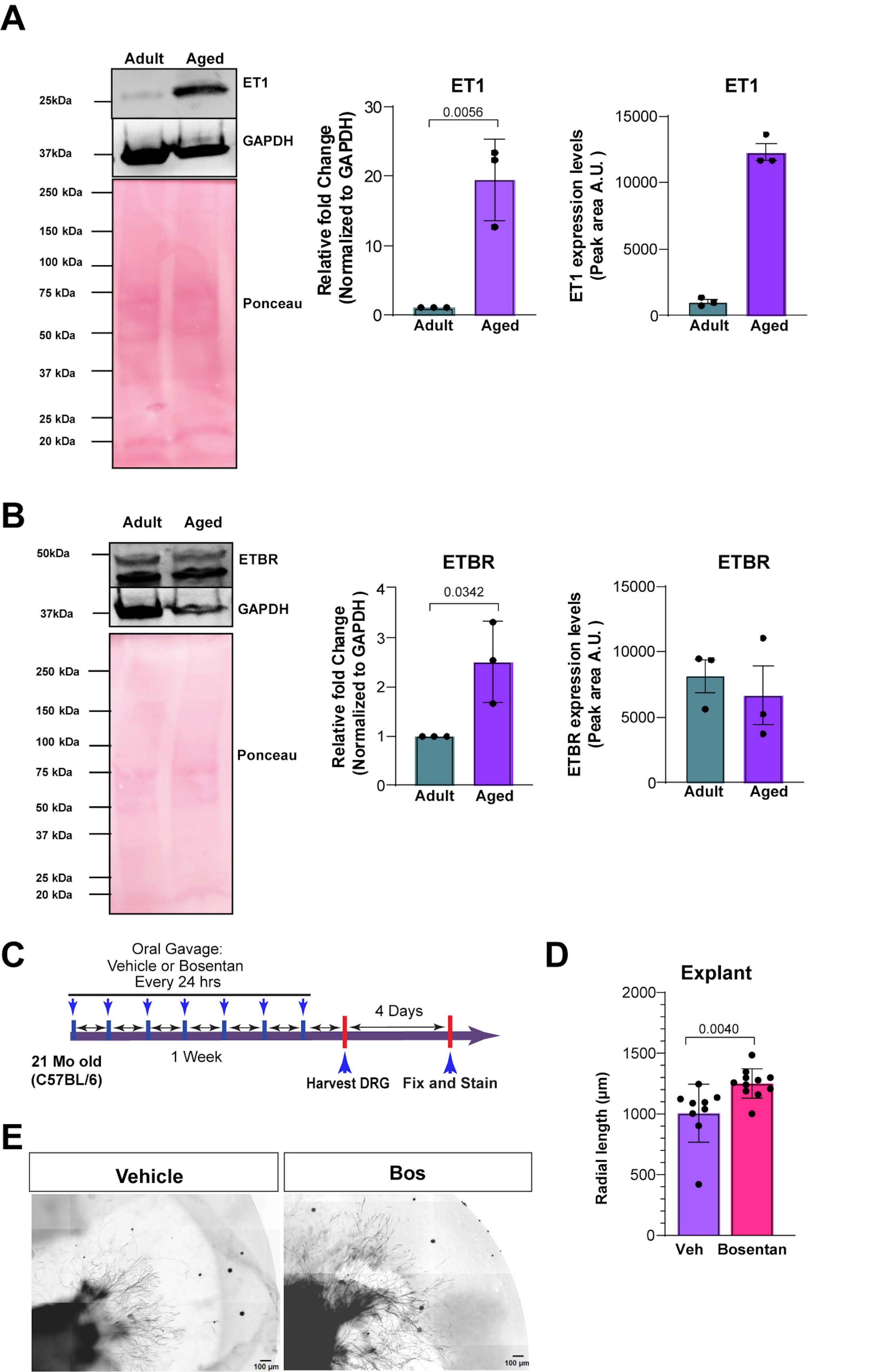

**Figure 5- Figure Supplement 1**

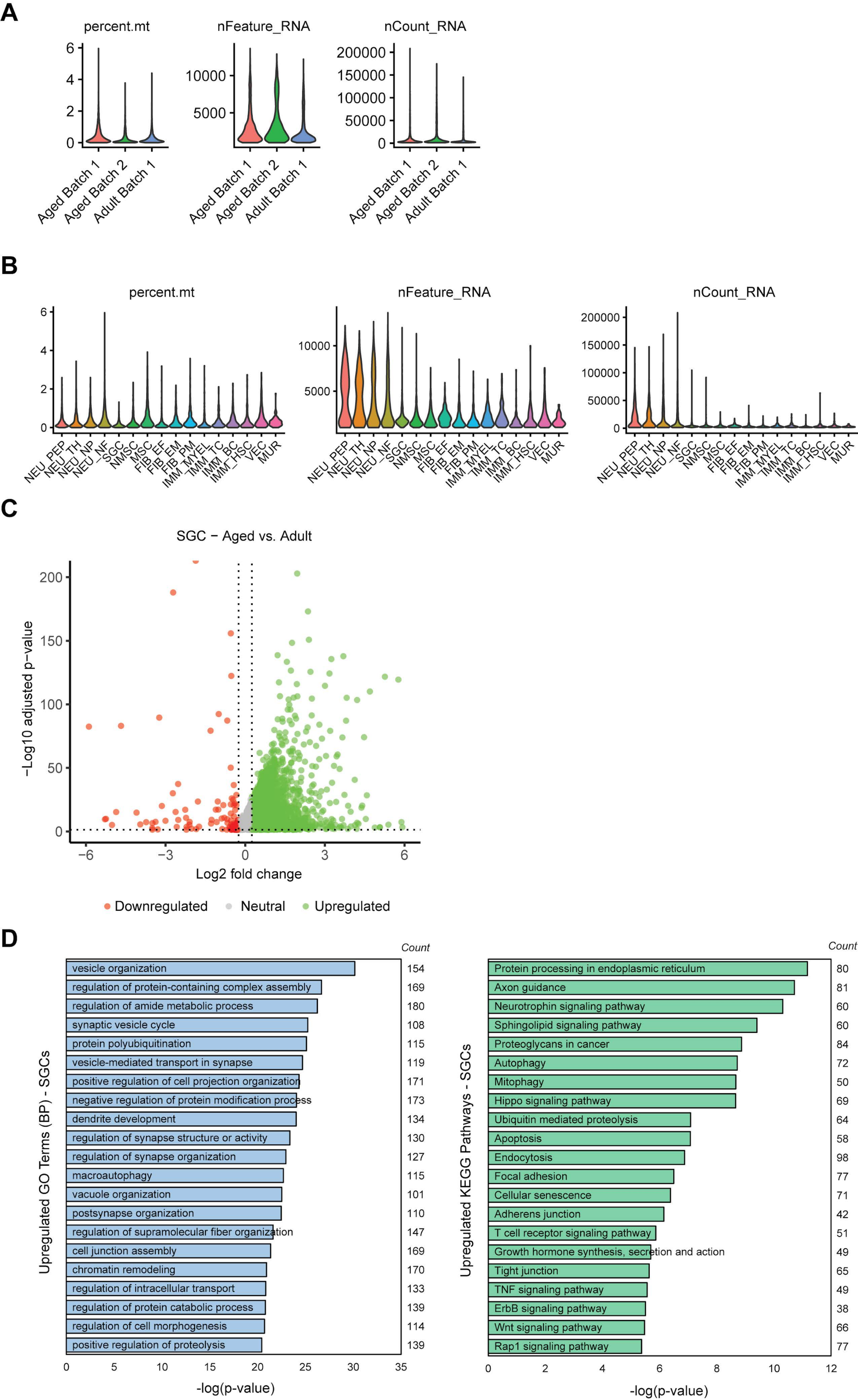

### Figure 6- Figure Supplement 1

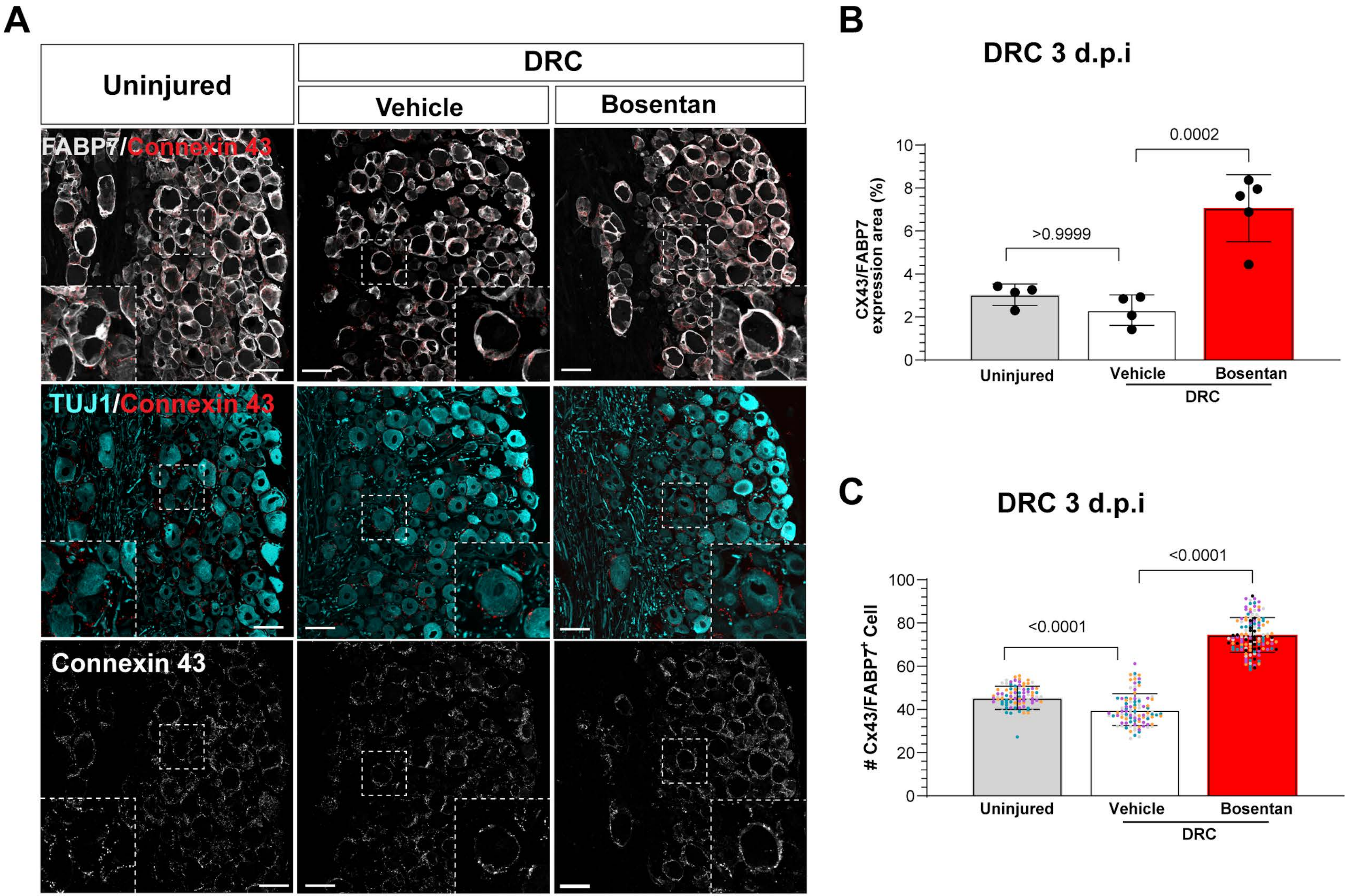
