## Supplementary material for "Endothelin B receptor inhibition rescues aging-dependent neuronal regenerative decline": Supp Table 1

**Supplementary Table 1**

| **Gene** | **Primer bank ID#** | **Sequence (5' -> 3')** | |
| --- | --- | --- | --- |
| *Gadph* | 6679937a1 | Forward Primer | AGGTCGGTGTGAACGGATTTG |
|  |  | Reverse Primer | TGTAGACCATGTAGTTGAGGTCA |
| *Atf3* | 31542154a1 | Forward Primer | GAGGATTTTGCTAACCTGACACC |
|  |  | Reverse Primer | TTGACGGTAACTGACTCCAGC |
| *Aif1* | 9506379a1 | Forward Primer | ATCAACAAGCAATTCCTCGATGA |
|  |  | Reverse Primer | CAGCATTCGCTTCAAGGACATA |
| *Fabp7* | 10946572a1 | Forward Primer | GGACACAATGCACATTCAAGAAC |
|  |  | Reverse Primer | CCGAACCACAGACTTACAGTTT |
| *Edn1* | 6753720a1 | Forward Primer | GCACCGGAGCTGAGAATGG |
|  |  | Reverse Primer | GTGGCAGAAGTAGACACACTC |
| *Ednra* | 14198449a1 | Forward Primer | ATGAGTATCTTTTGCCTTGCGG |
|  |  | Reverse Primer | GTCTTCCATGTGGCTGCTTAG |
| *Ednrb* | 6681269a1 | Forward Primer | GTGGCTTCTTGGGGGTATGG |
|  |  | Reverse Primer | TCTTAGTGGGTGGCGTCATTA |
